# Skin microbiome mirrors habitat divergence in amphibious combtooth blenny fish (Teleostei, Blenniidae)

**DOI:** 10.64898/2026.06.09.731066

**Authors:** Ewelina Rubin, Michele Felletti, Therese C. Miller, Bastian Bentlage, Diego. F. B. Vaz, Terry J. Ord, Iker Irisarri

## Abstract

Host–associated microbiomes play vital roles in organismal health, ecological interactions, and evolution, yet little is known about how microbial communities shift during the transition from aquatic to increasingly terrestrial habitats. Here, we characterize the skin microbiomes of three combtooth blenny species (*Blenniella paula*, *Praealticus labrovittatus*, and *Alticus arnoldorum*) that occupy distinct positions along the intertidal gradient in Guam—from fully subtidal (and exclusively aquatic) to intertidal (amphibious) and supratidal environments (exclusively terrestrial). Using 16S rRNA amplicon sequencing, we compared skin-associated bacterial communities with those in surrounding seawater and substrate biofilms to assess habitat influences on microbiome structure. Skin microbiomes were distinct from environmental microbial communities, indicating strong ecological filtering by the host. The divergence between skin and substrate microbiomes in the three species parallels their distribution along progressively higher zones of the intertidal gradient. The most divergent skin microbiome was that of the supratidal fish *A. arnoldorum*, characterized by higher Gammaproteobacteria abundance and enrichment of epiphytic and mucus-associated taxa. Across all species, we identified 32 microbial orders significantly enriched on the skin relative to environmental samples, including taxa commonly associated with fish mucosa (e.g., *Vibrio*, *Alteromonas*, *Cetobacterium*) and others rarely reported in aquatic marine fish (e.g., *Rubritalea*, *Granulosicoccus*). Several rare taxa with potential pathogenicity were also detected at low abundances. Together, these findings suggest that habitat-specific selective pressures strongly shape fish skin microbiomes along subtidal (aquatic) to supratidal (terrestrial) habitats and suggest that microbial symbionts may contribute to the ecological and physiological adaptations enabling amphibious lifestyles. This study provides the first comparative assessment of skin microbiome divergence across amphibious fish species along an intertidal gradient and offers a framework for predicting microbiome responses to environmental change.

## Introduction

Microbiome is increasingly recognized as an essential player in shaping the host’s function and fitness. Host and microbiome form the “holobiont”, which has been proposed as the unit of evolution upon which natural selection can act (Brucker and Bordenstein, 2012). In particular, skin microbiomes are essential for organisms’ overall health, survival, and adaptation to environmental conditions (Ross *et al*., 2019; Callewaert *et al*., 2020; Sehnal *et al*., 2021). The skin acts as a physical barrier that protects the body from harmful environmental factors and potential pathogens, and the skin-associated bacteria affect resistance and/or susceptibility to infections (Ross *et al*., 2019; Sehnal *et al*., 2021; Akat *et al*., 2022). In some fish species, the skin fulfills additional physiological roles, such as gas exchange (Glover *et al*., 2013). Skin microbiomes contain beneficial bacteria that enhance immunity and commensal bacteria that protect against pathogens through competitive exclusion (Llewellyn *et al*., 2014; Reverter *et al*., 2018; Sehnal *et al*., 2021). Often, fish skin is covered by a mucous layer that serves as a barrier and a site for microbial exchange, as it is in constant contact with the surrounding water (Esteban, 2012; Llewellyn *et al*., 2014). Thus, skin microbiomes are often more similar to the environment than internal gut microbiomes, the latter reflecting species-specific adaptations and being likely vertically inherited (Minich *et al*., 2022). Skin microbiomes may be used to study environmental perturbations, whereas gut microbiomes are more appropriate to identify phylosimbiosis—the correspondence between host phylogeny and microbiome (Sylvain *et al*., 2020).

While phylosymbiosis is crucial for skin microbiome composition, the environment plays an important role in microbiome shifts and modifications (Ross *et al*., 2019; Lim and Bordenstein, 2020; Bell *et al*., 2024). In aquatic animals, microbiomes are influenced by water salinity, temperature, and dissolved oxygen (Ross *et al*., 2019; Sehnal *et al*., 2021; McMurtrie *et al*., 2025). In commercially relevant fish, these have been extensively studied for their immediate implications in aquaculture (Gomez & Primm, 2021; Debnath *et al.,* 2023; Bell *et al.,* 2024; McMurtrie *et al.,* 2025), but analyses in wild settings that investigate ecological and evolutionary implications are less common, despite their importance for understanding host–microbe interactions.

One of the most extreme ecological shifts in fish is the conquest of intertidal and supratidal environments, or, more broadly, the transition from aquatic to terrestrial environments (Ord and Cooke, 2016). The importance of microbiomes in the process of terrestrialization has been proposed (Cannicci *et al*., 2020) but to date, only extremely fragmentary data exist, leaving this hypothesis largely untested. An analysis of gill-associated microbiomes in brachyuran crabs identified species-specific microbiomes, but the differences were not clearly linked to breathing adaptations (Bacci *et al*., 2023). Another study analyzed the microbiome of aestivating lungfish and identified granulocytes as key for pathogen defense (Heimroth *et al*., 2021). Aestivation, however, is a highly derived, extreme adaptation in some lungfish species that, besides adaptation to terrestrial stressors, it involves metabolic lethargy and cocoon secretion, additional aspects that can act as confounding factors in understanding the role of the microbiome in adaptation to out-of-water conditions.

Among fish, the family Blenniidae includes species with some of the most extreme amphibious adaptations to the supratidal environment (Ord and Cooke, 2016). Combtooth blennies are benthic fish that inhabit shallow coastal waters, including the intertidal and supratidal zones (Hundt *et al*., 2014). They lack scales and secrete thick mucus layers, two features that could have been adaptive for species that venture out of water. For example, they could facilitate cutaneous respiration and immune function in high-oxygen environments (Lemopoulos and Montoya-Burgos, 2021), provide protection against pathogens and desiccation, or reduce friction for locomotion out of water (Akat *et al.,* 2022; Esteban, 2012). Importantly, the lack of scales and the presence of a thick mucus layer could be associated with differences in the skin-associated microbiomes.

To date, there are no comparative studies of the skin microbiomes of combtooth blennies. However, three studies on the gut and gill microbiome of blennies provide some insight into the environmental drivers of their microbiomes. Hundt *et al*. (2017) analyzed the gut microbiota of two species with divergent diets: *Chasmodes saburrae*, which feeds on benthic or epiphytic crustaceans, and *Scartella cristata*, which feeds on plants and detritus. The increased abundance of *Vibrionaceae*, *Methylococcaceae*, and *Fusobacteriaceae* in *S. cristata* was linked to differences in their diets. Yoshida et al. (2022) compared the gut microbiomes of *Andamia tetradactylus*, a highly amphibious species feeding exclusively on algae, and *Entomacrodus stellifer*, which feeds on algae and invertebrates. This study also found critical diet-related differences, with a high prevalence of *Pseudomonadota* but an enrichment of *Spirochaetes* and *Tenericutes* in the gut microbiome of the strict herbivore. In addition, the absence of *Firmicutes* and *Actinobacteria* differentiated algae-eating blennies from the rest of the fish (Yoshida et al. 2022). Pratte et al. (2018) analyzed the gill and gut microbiomes of 15 reef fish families, including the blenny *Istiblennius* sp. Overall, the gill and gut microbiomes differed clearly within each species. However, microbiome similarity was higher between organs within the same species than between organs across different species. This pattern suggests that host-specific factors, potentially interacting with environmental exposure, play a stronger role than organ type alone in shaping bacterial communities in fish.

Despite extensive work on morphological and physiological adaptations (Hsieh, 2010; Egan et al., 2021), predation pressures (Ord et al., 2017), and abiotic conditions in rockpools (Ord et al., 2024), the role of the skin microbiome in the terrestrialization of combtooth blennies remains unexplored. Here, we study the skin microbiomes of three combtooth blenny species that occupy distinct habitats along the intertidal gradient in the Pacific island of Guam. *Blenniella paula* resides in the subtidal zone and water pools below the lowest tide line (Egan *et al*., 2021), representing the ancestral aquatic conditions of combtooth blennies (Ord and Cooke, 2016). *Praealticus labrovittatus* prefers the intertidal zone and is often found in rock pools above the tide waterline, as well as frequently out of the water entirely, and is regarded as amphibious (Platt *et al*., 2016). *Alticus arnoldorum* is considered exclusively terrestrial and inhabits rocks in the supratidal zone above the high tide line, which are constantly splashed but not submerged (Ord and Cooke, 2016; Platt *et al*., 2016). These three species exhibit various degrees of dependence on water, and thus could be considered as representatives of evolutionary “snapshots” of a hypothetical evolutionary path of increasingly amphibious lifestyles (Ord and Cooke, 2016).

The intertidal and supratidal zones are characterized by very variable and extreme conditions, including high water velocities and turbulence, fluctuating salinity and/or oxygen levels, and intermittent air exposure that can lead to desiccation; all factors that exert intense selective pressures on fish morphology and physiology (Hsieh, 2010; Egan *et al.,* 2021). By contrast, subtidal habitats offer greater stability and structural complexity, allowing for niche partitioning and ecological specialization that may be reflected in skin microbiome compositions (Chiarello *et al*., 2018). Thus, we hypothesized that the skin microbiome of species inhabiting subtidal, intertidal, and supratidal habitats should reflect adaptations to these divergent environments. This is the first study to examine microbiome shifts in species occupying habitats along the water–land gradient, providing new insight into selective pressures on skin microbiomes associated with amphibious behaviors. We found marked differences along the intertidal gradient, providing a new perspective on the relationship between habitat and skin microbiome composition and fish adaptation to terrestrial conditions. These findings have broader implications for understanding evolutionary transitions from aquatic to terrestrial life, suggesting that microbial communities may have played a role in such adaptations. Moreover, by linking habitat characteristics to microbiome composition, this research offers a framework for predicting microbiome responses to future environmental changes in coastal ecosystems.

## Materials and methods

### Sampling

Ten individuals each of *A. arnoldorum, P. labrovittatus,* and *B. paula,* were caught in Pago Bay, Guam, on May 18 and 19, 2023. Individuals were caught in the intertidal zone using hand-held nets and transferred into new zip-lock bags, each containing 50 mL (*A. arnoldorum* and *P. labrovittatus*) or 100 mL (*B. paula*) of autoclaved seawater. Fish were left in the bags for 30 seconds to rinse off external surfaces that could have been contaminated during fishing and handling. Water was removed, and fish were swabbed (both ventral and dorsal areas) to sample mucus microbiomes. To assess the relative contribution of the environment to skin microbiomes, we sampled the biofilms on the substrates where the three species were found: subtidal rock holes for *B. paula*, rock pools for *P. labrovittatus*, and supratidal rocks for *A. arnoldorum*. In the case of *B. paula*, we sampled the exact substrate on which each fish was residing; thus, the number of fish skin samples equals the number of fish substrates. For the other two fish species, which are more mobile, we sampled rocks or rockpools where the fish were found, but the numbers of skin and substrate do not exactly correspond. For comparison, we also collected 3 L of seawater samples from our general sampling area. Specifically, we sampled nearshore surface water near the *A. arnoldorum* and *P. labrovittattus* habitat and again approximately 10 m from shore, corresponding to the *B. paula* habitat. All seawater samples were collected in triplicate on different days: May 18 and May 20, 2023. Seawater samples were filtered using a vacuum pump (GAST) through either a 1.2 μm or 0.45 μm pore filter (Whatman). Fish skin swabs and filters were placed into 2 mL tubes and stored at −80 °C until further processing.

### Microbiome profiles

DNA was extracted using the ZymoBIOMICS DNA extraction kit (Zymo Research) following the manufacturer’s protocol. The V4 region of the 16S rRNA gene was amplified using modified forward 515F (Parada *et al*., 2016) and reverse 806R (Apprill *et al*., 2015) primers. The product from the first PCR was used in the second PCR, which was designed to complete the addition of Illumina adapters and the 6-mer dual indices as previously described by (Kozich *et al*., 2013). Both PCRs were generated using Takara PCR buffer and Hot Start polymerase (Takara). The first PCR was carried out in 30-μL reactions with the following final concentrations: dNTPs at 0.8 mM, each primer at 0.5 µM, and 30-100 ng of total DNA. The first PCR cycling conditions were as follows: a hot-start activation step for 2 min at 95 °C, followed by 30 cycles of 95 °C for 40 s, 58 °C for 2 min, and 72 °C for 1 min, with a final extension of 5 min at 72 °C. The PCR was purified using GeneJet (ThermoFisher). The second PCR was performed in a 20-µL reaction volume using the same components and final concentrations as above, with 30 ng of the first PCR product serving as the template. The second PCR cycling conditions were as follows: a hot-start activation at 95 °C for 2 min, followed by five cycles of 95 °C for 40 s, 59 °C for 2 min, and 72 °C for 1 min, and concluded with a final extension at 72 °C for 7 min. All samples were pooled and gel-purified using the Qiagen gel purification kit. The V4-16S library pool was sent to Michigan State University Research Technology Support Facility Genomics Core for Illumina MiSeq sequencing, using a 2 x 250 bp cycling kit.

Data quality of the raw sequencing reads for each sample was evaluated using the FastQC tool version 0.12.1 (Brown *et al*., 2017). The remaining analyses were conducted in R v. 4.5.1 using multiple packages, including dada2 v. 1.34 (Callahan et al. 2016), phyloseq (McMurdie and Holmes 2013), microbiome (Leo Lahti 2017), and microViz (Barnett D.J.M.). Specifically, after quality inspection, the raw reads were trimmed and filtered using the dada2 package with a minimum Phred quality score of 10. The same package was used to generate the ASV (Amplicon Sequences Variants) sequence table and to remove chimeric ASVs. Taxonomy was assigned using the SILVA SSU rRNA database v.138.2 (Quast *et al*., 2013). Following common practice in bacteriology, all scientific names, including those above the genus level, are highlighted in italics (Thines et al. 2020). ASVs classified as mitochondria, chloroplasts, eukaryotes, or an unknown kingdom were removed from the final count table. Count data were transformed to relative abundance and visualized using stacked bar charts constructed in the library fantaxtic (Teunisse, 2022). All other plots were constructed using the ggplot2 (Wickham, 2016). Differential abundance (DA) analysis was performed using the corncob R package (Martin et al., 2020). Functional profiles of all ASVs derived from the fish skin samples (2,670 ASVs) were inferred using PICRUSt2 v2.5.2 (Douglas *et al*., 2020) with default parameters. Predicted KEGG Ortholog (KO) abundances were generated for each ASV and subsequently merged with taxonomic assignments obtained from the SILVA database as determined above. KEGG Orthologs were mapped to the KEGG BRITE hierarchy using the three-tier classification system (levels A–C; (Kanehisa and Goto, 2000). To quantify the relative contribution of bacterial genera to predicted metabolic functions, KO abundances were aggregated by genus and KEGG BRITE level C category by summing normalized taxon–function contributions (norm_taxon_function_contrib). For each KEGG level C category, the proportional contribution of each genus was calculated as a percentage of the total predicted functional abundance for that category. These genus-level functional contributions were visualized using heatmaps generated by the ComplexHeatmap R package (Gu *et al*., 2016). As PICRUSt2 provides predictions based on closest matching reference genomes, functional profiles should be interpreted as putative rather than experimentally validated metabolic capacities. All other R code for all full analyses and for generating Figures is available in the GitHub repository: https://github.com/ewrubin/combtooth-blenny-microbiome

## Results

### Divergence of skin microbial profiles reflects increasingly terrestrial lifestyles

In this study, we generated bacterial microbial profiles for the skin of three combtooth blenny fish species that inhabit increasingly higher zones of the intertidal gradient: *B. paula* in the subtidal areas, *P. labrovittatus* in intertidal rockpools, and *A. arnoldorum* in supratidal rocks. The skin microbial profiles of all three species were markedly different from the microbial communities found in their surrounding environment, whether in seawater or on the substrate they inhabited (Figure 1A; PERMANOVA, F = 7.37, p < 0.01). However, beta-dispersion analyses also detected significant differences in within-group variability among environments (Figure 1A, permutest, F = 22.51, p = 0.001), with seawater communities exhibiting lower dispersion than both fish skin and substrate-associated communities. Among environmental samples, seawater bacterioplankton was the most distinct, clearly separating from both the skin and rock biofilm microbial profiles (Figure 1A). Differences between skin and substrate microbiomes increased with the increasing terrestriality of the fish species. They were most similar in the subtidal *B. paula*, with overlap observed in two cases, and increasingly divergent in the intertidal *P. labrovittatus* and the supratidal *A. arnoldorum* (Figure S1). Skin microbiomes of the three fish species were species-specific (Figure 1B; PERMANOVA, F = 5.79, p < 0.01), with the supratidal *A. arnoldorum* displaying the most dissimilar profile (Figure 1B). No significant differences in beta dispersion were detected among host skin microbiomes (Figure 1B, permutest, F = 2.98, p = 0.07), suggesting that the observed PERMANOVA result primarily reflected differences in community composition rather than within-group variability. In addition, substrate-associated bacterial communities exhibited substantial overlap among host species in ordination space (Figure 1B), but PERMANOVA detected a significant effect of species identity on community composition (F = 2.85, p = 0.001). Again, the beta-dispersion analyses also indicated significant differences in within-group dispersion (permutest, F = 3.97, p = 0.04), suggesting that the observed PERMANOVA result may partially reflect differences in community variability among host-associated substrates.

**Figure 1.**
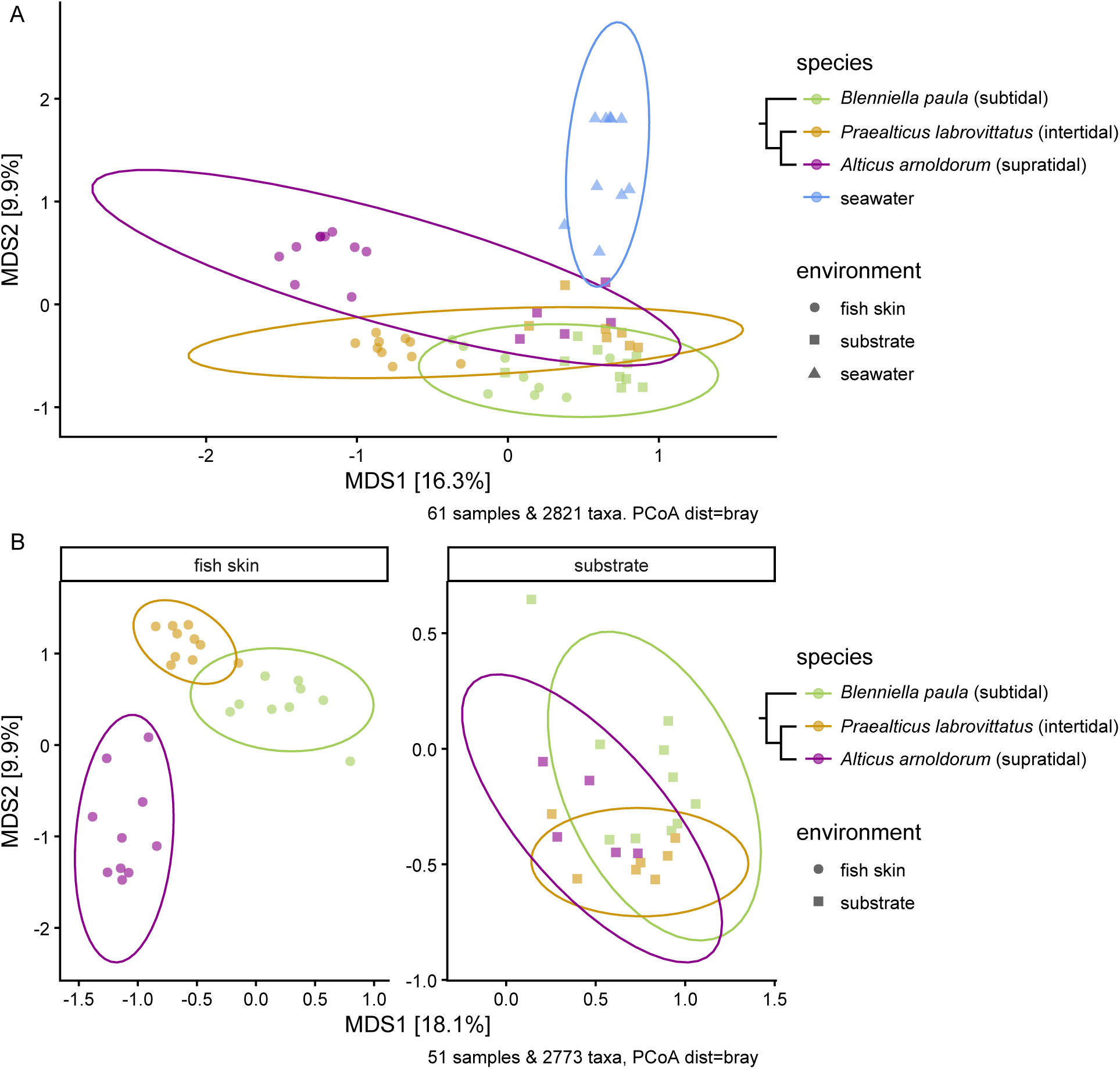
Principal coordinates analysis (PCoA) based on Bray–Curtis dissimilarities of bacterial communities associated with seawater, fish skin, and rock substrate samples. (A) Ordination of all samples. (B) Fish skin and substrate biofilm communities associated with the three combtooth blenny species. Ellipses indicate 95% confidence intervals around group centroids. Phylogenetic relationships and habitat of host fish are indicated.

Alpha diversity indices showed that microbial communities were more diverse in seawater and substrate than on fish skin (Figure 2; Kruskal–Wallis, H = 31.105, p < 0.01). The Shannon diversity index (mean ± SD) was 5.65 ± 0.417 for seawater, 5.75 ± 0.552 for substrate, and 4.47 ± 0.682 for fish skin (Figure 2). The difference in alpha diversity was significant between fish skin and seawater and between fish skin and substrate (Wilcoxon test, p < 0.01), but not between seawater and substrate. Skin microbiomes of the three fish species did not differ significantly in alpha diversity.

**Figure 2.**
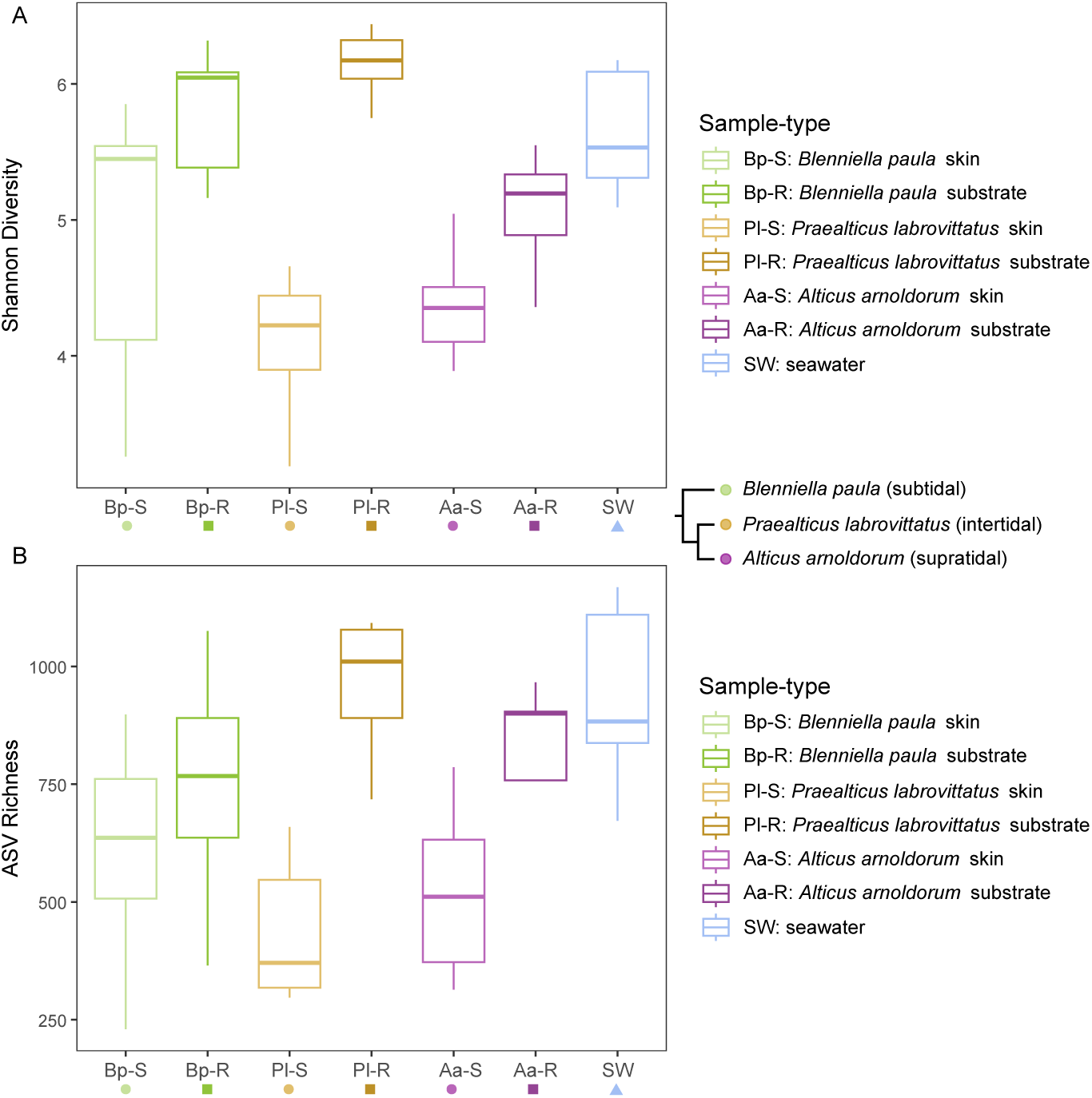
Alpha diversity indices of microbial communities from fish skin, rock substrate, and seawater samples. Shannon diversity (A) and ASV richness (B) are shown for skin (S) and substrate (R) microbiomes associated with *Blenniella paula* (Bp), *Praealticus labrovittatus* (Pl), and *Alticus arnoldorum* (Aa), and seawater (SW).

### Differential bacterial abundance at the class, order, and genus levels

*Gammaproteobacteria* were the most abundant class in fish skin microbiomes, accounting for 19%-68% of total bacterial ASVs (Figure 3). Their abundance was significantly higher on the fish skin than on either seawater or substrate (Figure 3, Table S1, Wald test, FDR < 0.05). In addition, among other classes with increased abundances on fish skin in comparison to either seawater or substrate were *Fusobacteriia*, *Verrucomicrobiia*, and *Bacilli*, contributing between 7-34%, 3-11%, and 2-5% to the total abundance, respectively (Figure 3, Table S1, Wald test, FDR < 0.05). On the other hand, *Bacteroidia, Cyanobacteriia, Desulfobulbia,* and *Deinococci* were more abundant in seawater and substrate in comparison to fish skins (Figure 3, Table S1, Wald test, FDR < 0.05). Classes with an average relative abundance below 0.3 % across the dataset are grouped into “other” in Figure 3; therefore, not all differentially abundant classes are displayed. *Alphaproteobacteria* were also abundant across all sample types, contributing between 3% and 43% to the fish skin microbiome, between 19% and 43% to seawater, and between 16% and 31% to the substrate. Unlike other bacterial groups, their abundance did not differ significantly among the three environments (Figure 3, Wald test, FDR > 0.05) but was differentially abundant among hosts, with lower abundance in *A. arnoldorum* (7%) compared to the other two species (23% in both) (Table S2, Wald test, FDR < 0.05).

**Figure 3.**
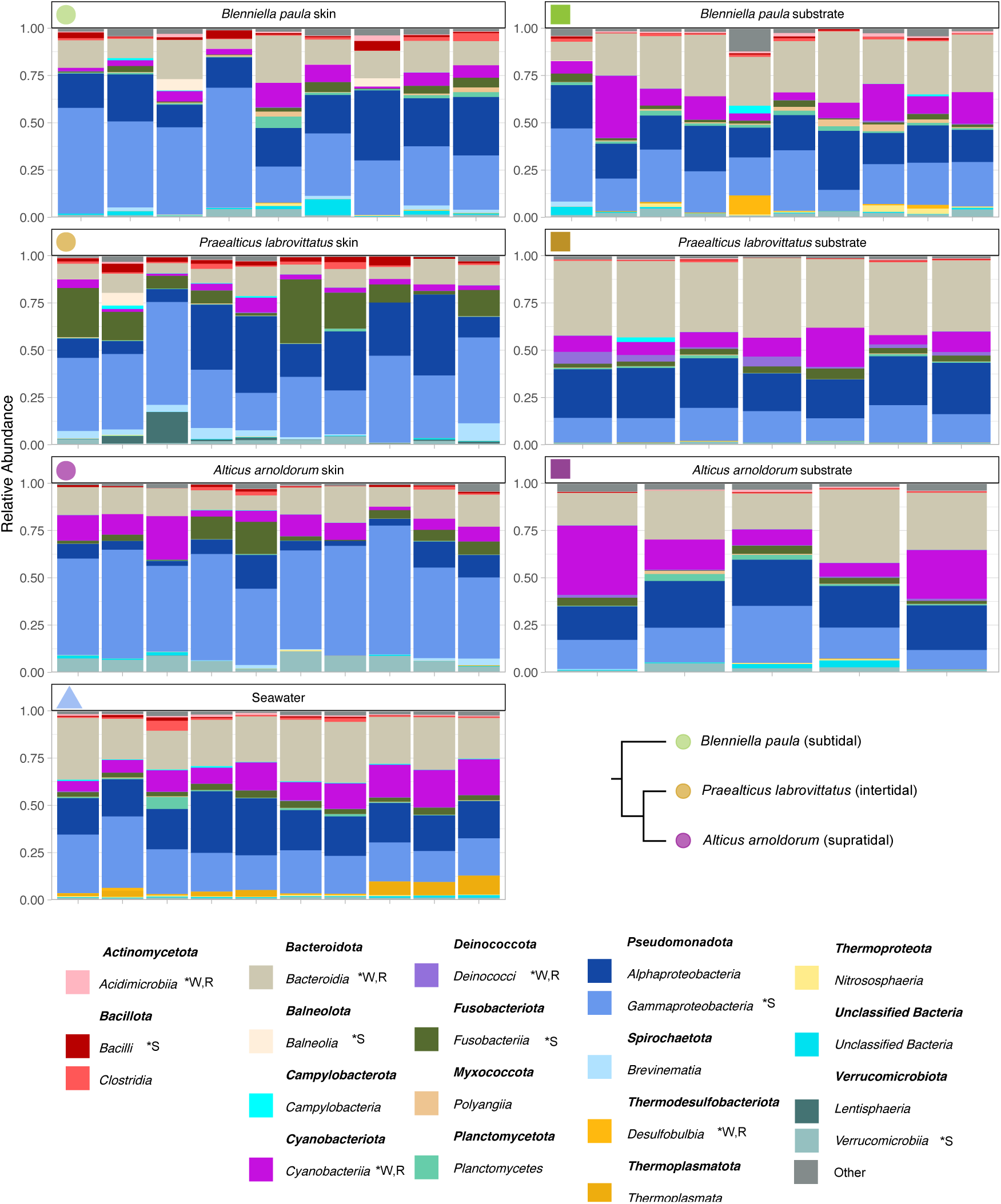
Relative abundance of the 20 most abundant microbial classes across fish skin microbiomes, associated rock substrate biofilms, and seawater samples from the three studied host species. Letter designations indicate classes that were significantly enriched either in fish skin relative to seawater or substrate (*S), or in both seawater and substrate relative to fish skin (*W, R), as determined by beta-binomial regression and Wald tests (FDR < 0.01).

Out of the 128 order-level groupings, 79 (57%) were differentially abundant among the three environments: seawater, substrate, and fish skin (Table S3; Wald test, FDR < 0.05). Of these, 16 were more abundant on fish skin than in either the seawater or substrate microbiomes (Table S2, Wald test, FDR < 0.05), with the most substantial increases for *Kordiimonadales*, *Cardiobacteriales*, *Spirochaetales,* and *Burkholderiales* (Table S3). Out of the 79 orders, 14 were more abundant in seawater and substrate compared to fish skin (Table S3; Wald test, FDR < 0.05). Regarding differences among skin microbiomes of the three fish species, 43 orders were differentially abundant (Table S4; Wald test, FDR < 0.05). Six microbial orders were enriched in the supratidal *A. arnoldorum* compared to both *B. paula* and *P. labrovittatus,* including *Granulosicoccales*, *Cardiobacteriales*, *Arenicellales*, *Verrucomicrobiales*, and *Phormidesmiales* (Table S4, Wald test, FDR < 0.05). By contrast, *Kordiimonadales*, *Pseudomonadales*, and *Balneolales* and several other orders were more abundant on the skin of *B. paula* and *P. labrovittatus* than on *A. arnoldorum* (Table S4; Wald test, FDR < 0.05).

The top-most abundant genera (29 total) contributing more than 1% to the total abundance of fish skin microbiomes for any host species are shown in Figure 4. The dominant genera were *Vibrio* (average abundance of 7.1%)*, Cetobacterium* (5.5%), and *Alteromonas* (4.9%) (Figure 4). Eighteen most frequent genera were differentially abundant among the three host species (Figure 4A, Wald test, FDR < 0.05), and an additional 11 most frequent genera had not different relative abundance between the three host species (Figure 4B). Eight of the top genera (*Arenicella*, *Gammaproteobacteria UG*, *Granulosicoccus*, *Lewinella*, *Maribacter*, *Phormidesmis*, *Rubritalea, Schizothrix)* were enriched in *A. arnoldorum*; two (*Cetobacterium* and *Kordiimonas*) in *P. labrovittatus;* and one (*Acinetobacter*) in *B. paula* (Figure 4, Wald test, FDR < 0.05). And the undescribed genus of *Burkholderiales UG* was enriched on the skin of both *B. paula* and *A. arnoldorum* (Figure 4, Wald test, FDR < 0.05). Additional lower abundance genera were also differentially abundant among the host species (Table S5, Wald test, FDR < 0.05).

**Figure 4.**
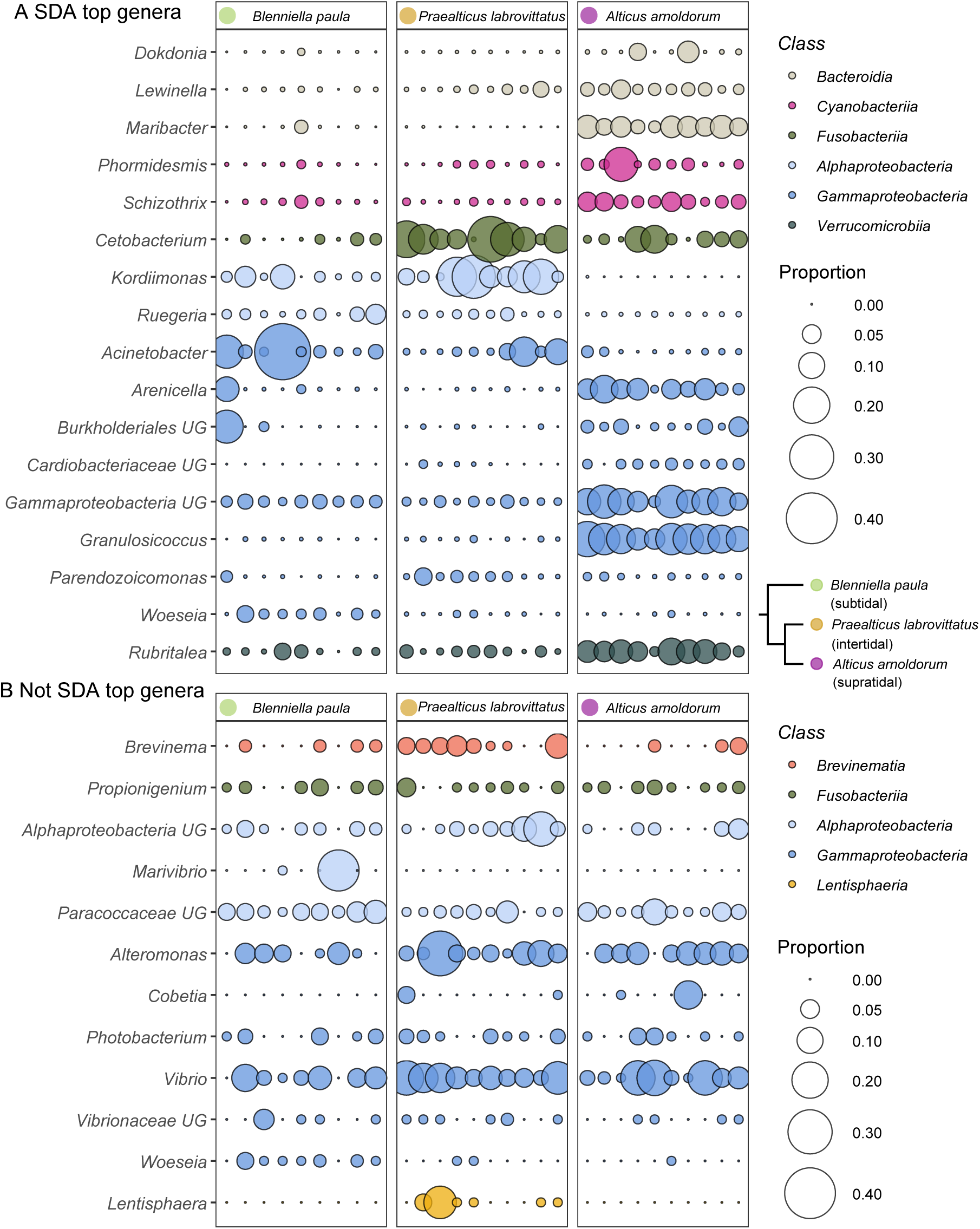
Most abundant bacterial classes (mean relative abundance ≥1% in at least one host species), colored by class and separated into (A) statistically differentially abundant (SDA) genera (Wald test, FDR < 0.05) and (B) non-differentially abundant genera across the three host species. UG: undescribed genus.

### Functional potential of bacteria associated with fish host skins

The relative contribution of microbial taxa to functional potential was assessed by calculating the proportional contribution of all 552 bacterial genera detected on fish skin to 140 KEGG level C metabolic categories (Table S6). For visualization, we focused on genera that were ecologically meaningful contributors by selecting those that reached at least 1% relative abundance in the skin microbiome of at least one host species, including those eighteen that were differentially abundant among the host species (Figure 4A, Table S6) and those that did not differ in relative abundance (Figure 4B). Several metabolic categories were dominated by a small number of differentially abundant genera, revealing strong taxon-specific functional signatures across host species. Of particular interest were *Cetobacterium*, *Kordiimonas*, and *Parendozoicomonas*, which were enriched on the skin of *P. labrovittatus*. *Cetobacterium* contributed most to xenobiotic degradation pathways, accounting, among others, for ca. 15% to nitrotoluene degradation. *Kordiimonas* contributed over 11% to the degradation of pinene, camphor, and geraniol, and more than 8% to xenobiotic metabolism via cytochrome P450. *Parendozoicomonas* contributed more moderately but consistently across several metabolic categories associated with xenobiotic and secondary metabolism (Figure 5).

**Figure 5.**
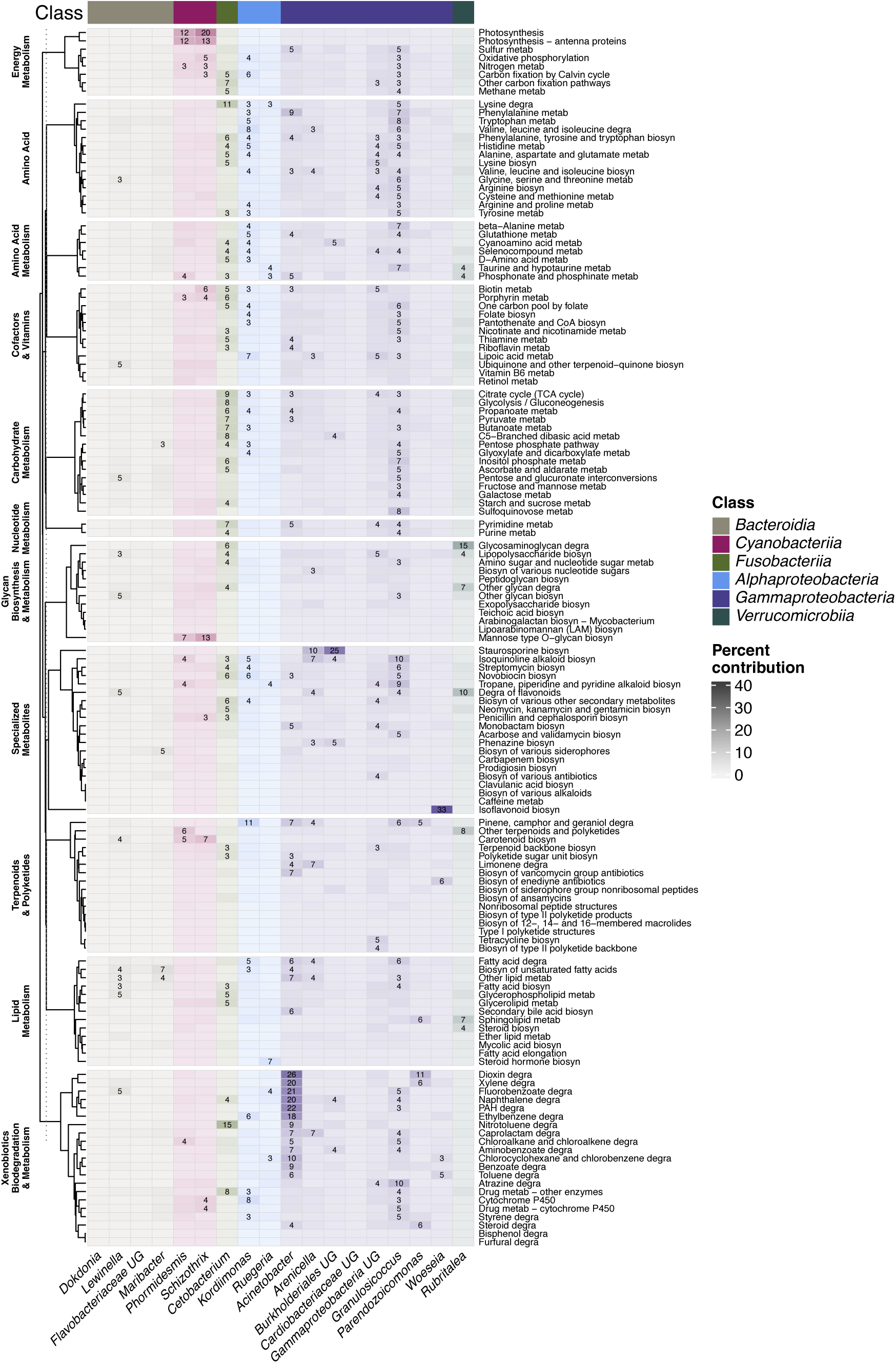
Predicted functional metabolic profiles of the most abundant bacterial genera associated with fish skin microbiomes. Genera shown were statistically differentially abundant among the three host species (Wald test, FDR < 0.05) and contributed a mean relative abundance ≥1% in at least one host species. Genera are grouped and color-coded by bacterial class. Functional profiles were inferred using PICRUSt2 and are shown as the percentage contribution of each genus to individual KEGG level-C pathways. UG: undescribed genus.

The skin microbiome of *B. paula* was characterized by strong functional contributions from *Acinetobacter* and *Woeseia*. *Acinetobacter* was the dominant contributor to multiple xenobiotic biodegradation pathways, accounting for 26% of dioxin, 21% of polycyclic aromatic hydrocarbons, and approximately 20% of fluorobenzoate, naphthalene, and xylene. *Woeseia* contributed significantly to specialized metabolite biosynthesis, and 14% of enediyne antibiotic biosynthesis (Figure 5).

Energy metabolism was largely driven by cyanobacteria enriched on the skin of *A. arnoldorum.* Specifically, *Phormidesmis* and *Schizothrix* were the primary contributors to photosynthesis-related pathways, including photosynthesis and photosynthesis-antenna protein synthesis (Figure 5). In addition, *Granulosicoccus* and *Rubritalea*, also enriched on *A. arnoldorum,* contributed substantially to 15% of glycosaminoglycan metabolism and ca. 10% of flavonoid degradation. Several other genera contributed disproportionately to specialized metabolite pathways. An undescribed genus within *Burkholderiales* accounted for ca. 25% of staurosporine biosynthesis and *Arenicella* contributed ca. 10% to the same pathway (Figure 5). These contributions highlight the role of less abundant but functionally specialized taxa in shaping the metabolic landscape of the skin microbiome.

Several of the most abundant genera were not differentially abundant among host species but made substantial functional contributions. For example, *Vibrio* contributed ca. 10% to sulfoquinovose metabolism, while *Alteromonas* contributed ca. 10% to flavonoid degradation. Additional contributions included 12% to galactose metabolism and 11% to glycan degradation by shared core taxa. A genus-level taxon within *Paracoccaceae* (139 ASVs and 46 reference genomes) also contributed prominently to xenobiotic degradation, accounting for 14% of chlorocyclohexane and chlorobenzene degradation, 12% of atrazine degradation, and 10% of caprolactam degradation (Figure S2).

## Discussion

### Fish skin as an ecological filter on environmental microbiomes

The skin microbiomes were significantly different and less diverse than the bacterioplankton in seawater and the biofilms on the substrates on which fish lived. This suggests that fish skin acts as an ecological filter for the bacterial species available in the environment. Our findings contrast with those of Pratte *et al*. (2018), which found higher alpha diversity in the gill and gut microbiomes of fish than in the environment. However, the species, geographical location, and tissues in Pratte *et. al*. (2018) differed from ours. The ecological filter is further corroborated by the marked differences between fish skin microbiomes and the substrates on which they live. This pattern was consistent across all three host fish studied. Moreover, skin microbiomes increasingly diverged among species inhabiting increasingly higher areas along the intertidal gradient, while substrate microbiomes were largely overlapping. Substrate and skin microbiomes were most similar for the subtidal *B. paula,* which lives in a tight association with the substrate in rock holes and crevices in the reef flat. Substrate and skin microbiomes diverged for *P. labrovittatus,* which inhabits intertidal rockpools, and were most dissimilar between substrate and skin of *A. arnoldorum,* which typically inhabits supratidal rocks (Myers, 1999). This suggests that the intertidal gradient imposes a second ecological filter on the bacterial species that occur on fish skin, a filter that increases with the increasingly terrestrial lifestyles of fish hosts.

### Habitat shapes skin microbiome in combtooth blennies

The three fish species inhabiting the subtidal, intertidal, and supratidal zones diverged markedly in their skin microbiomes, mirroring their level of terrestriality. Host-specific profiles indicate that skin microbiomes are largely shaped by intrinsic factors, in agreement with previous research (Gomez and Primm, 2021; Sehnal *et al*., 2021). Microbiomes often display a phylosymbiotic signal, meaning that microbial communities reflect the evolutionary divergence of host species. For example, skin microbiomes were more similar within rather than between fish host families in a study of seven wild fish species of four families (Labridae, Sebastidae, Sparidae, and Rajidae) off the southern coast of Korea (Han *et al*., 2024). Other studies found that environmental factors such as habitat and diet can significantly alter the phylosymbiotic signal (Bell *et al*., 2024). In our study, the skin microbiome profiles of *B. paula* and *P. labrovittatus* were more similar to each other than those of *A. arnoldorum*. This pattern does not reflect phylogenetic affinities, as *P. labrovittatus* is more closely related to *A. arnoldorum* than to *B. paula* (Figure 1; Hundt et al. 2014; Muñoz-Sánchez, et al. 2026). Instead, skin microbiomes reflect habitat similarity in the intertidal ecotone: *B. paula* and *P. labrovittatus* remain largely submerged, respectively, in the subtidal and in intertidal rock pools, whereas *A. arnoldorum* has a terrestrial lifestyle on rocks above the waterline, consistent with its more divergent microbiome profile. This suggests that environmental conditions on rocks above the water line are different enough from those in the subtidal or rockpools to override potential phylosymbiotic signal in the microbiome composition. Adult *A. arnoldorum* live out of water and their activity typically peaks around mid-tide and with moderate temperatures, individuals retreating to shelter in rock holes and crevices at low tides (Ord and Tonia Hsieh 2011; Ord and Cooke 2016). Thus, *A. arnoldorum* likely experiences markedly different conditions in the supratidal, with higher light and UV exposure, more variable temperatures, increased osmotic stress and desiccation risk, and distinct food resources and pathogen communities. Differences between the skin microbiomes of *B. paula* and *P. labrovittatus* are likely associated with particular environmental conditions in their preferred habitats. Unlike subtidal waters, tidal pools like those inhabited by *P. labrovittatus* can have large fluctuations in temperature and dissolved oxygen and other abiotic factors (Ord *et al*., 2024), which could affect the skin microbiome composition.

### Gammaproteobacteria and Alphaproteobacteria dominate fish skin microbiomes

This study identified six bacterial classes and 16 orders that were more abundant on the skin of the three blenny species than in seawater or the substrate where they lived. Among these, *Gammaproteobacteria* were the most abundant across all three species (Figure 2), with a statistically higher relative abundance in *A. arnoldorum* than in the other two hosts. This class also showed statistically higher relative abundances in skin (43%-68%) than in substrate (10-39%) and seawater (16-38%). *Gammaproteobacteria* are the dominant group in the skin microbiomes of many marine fish (Larsen *et al*., 2013; Gomez and Primm, 2021; Bell *et al*., 2024). The relative abundance of *Alphaproteobacteria* was similar in seawater (19-33%), substrate (16-31%), and fish skin (3-43%), but varied significantly among hosts, with lower abundance in *A. arnoldorum* (7%) than in the other two species (23% in both). Thus, the skin microbiome of the highly amphibious *A. arnoldorum* is characterized by higher proportions of *Gammaproteobacteria* and lower proportions *of Alphaproteobacteria*.

Within *Gammaproteobacteria*, fish skins were enriched in bacterial orders *Enterobacterales, Burkholderiales, Francisellales, Arenicellales, Cardiobacteriales*, and *Granulosicoccales*. Members of the first three orders have been reported in fish microbiomes (Debnath *et al*., 2023; Bell *et al*., 2024; Jacobsen *et al*., 2025) whereas the latter three have not and may be unique to our species or location. Notably, *Granulosicoccus*, a species found in Antarctic seawater (Baek *et al*., 2014) or as epiphytes on marine algae (Kurilenko *et al*., 2010; Park *et al*., 2014), was among the top five most abundant genera across all three species and was especially enriched in *A. arnoldorum*. The predicted functional profile of *Granulosicoccus* revealed that it contributes the highest proportion to the degradation of atrazine (10%) and sulfoquinovose (8%). Atrazine is the most widely used herbicide globally and reaches coastal waters through runoff, impacting mostly photosynthetic and herbivorous organisms (Graymore *et al*., 2001) whereas sulfoquinovose is a sulfated monosaccharide produced by photosynthetic organisms and a source of carbon and sulfur for many environmental bacteria such as *Granulosicoccus*.

The dominant *Enterobacterales* (2.8–48.5%) genera were *Vibrio* (0.4–19%), *Alteromonas* (0.2–31%), and *Photobacterium* (0.1–3.6%), whose abundances did not differ among fish hosts. These genera commonly dominate the microbiomes of marine fish skin, yet their functional roles remain poorly understood (Chiarello *et al*., 2018; Bell *et al*., 2024; Han *et al*., 2024; Jacobsen *et al*., 2025). Recently, Yoshida et al. (2022) found a high prevalence of *Vibrio* species in the guts of another two combtooth blenny species. They isolated and sequenced the genomes of two *Vibrio* strains that encode multiple digestive enzymes, including alginate lyases, amylases, chitinases, and mannosidases, which could help degrade algal polysaccharides in fish guts. In our study, *Vibrio’s* highest functional contributions were to sulfoquinovose metabolism, whereas *Photobacterium’s* were to glycan degradation and galactose metabolism, and *Alteromonas’* to flavonoid degradation. Thus, in agreement with Yoshida et al. (2022), we found that a consistent portion of *Enterobacteriales* likely catabolize specifically compounds such as sulfoquinovose and flavonoids produced by algae and cyanobacteria or, in the case of *Photobacterium,* glycosaminoglycans (GAGs) from the fish skin mucus (Scholz *et al*., 2021; Fernando *et al*., 2022; Chen *et al*., 2026). A key limitation of our barcoding approach is that 16S-V4 lacks resolution to distinguish among species or strains of *Vibrio* and other *Enterobacteriales*. This is important because some *Vibrio* strains are symbiotic and others are pathogenic and can cause skin lesions under stress (Manchanayake *et al*., 2023). Similarly, some *Photobacterium* species are pathogenic whereas others are mutualistic or bioluminescent (Labella *et al*., 2017; Moi *et al*., 2017). Strain-level resolution and experimental testing are therefore necessary to clarify the functions of these bacterial genera in greater detail. Nonetheless, their consistent high abundance in healthy fish suggests that many species are commensal or beneficial, but we cannot rule out the possibility of them being opportunistic pathogens.

Additional *Gammaproteobacteria* genera with predicted interesting functional profiles include *Arenicella* (an undescribed genus –UG– within the *Burkholderiales*), *Acinetobacter,* and *Woeseia*. *Arenicella* had a higher relative abundance in the *A. arnoldorum* skin microbiome, and *Burkholdariales UG* was enriched in *A. arnoldorum* and *B. paula* compared to *P. labrovittatus*. Both *Arenicella* and *Burkholderiales UG* were predicted to contribute significant proportions to the staurosporine biosynthesis KEGG module (10% and 25%, respectively), which comprises a number of tryptophane-derived antimicrobial and antifungal specialized metabolites. Manual examination of this pathway revealed the presence of three enzymes required for pyrrolnitrin production (Table S7), an antifungal compound produced by some symbiotic bacteria that can protect their host from fungal infection (Hammer *et al*., 1999; Hwang *et al*., 2002). A metagenomic analysis might confirm the presence of this function in the microbiome of these fish. *Acinetobacter* had the highest relative abundance on the skin of the subtidal *B. paula* in comparison to *P. labrovittatus* and *A. arnoldorum.* Its predicted functional role is to protect the host from persistent marine pollutants, including polycyclic aromatic hydrocarbons (PAHs), which tend to accumulate in marine sediments (Ghosal *et al*., 2016) and in some Guam locations can be found above the US national average (Denton *et al*., 2006).

*Alphaproteobacteria* found at higher relative abundances in the skins of submerged blennies are *Kiloniellales*, *Kordiimonadales*, and *Rhodospirillales*. Of these, *Kordiimonas* and *Ruegeria* were most prevalent in submerged *B. paula* and P*. labrovitattus* than in the supratidal *A. arnoldorum*. *Kordiimonas* was predicted to be important for the degradation of specialized metabolites (terpenoids) and the removal of xenobiotics via cytochrome P450 activity, (Greule *et al*., 2018; Behrendorff, 2021). *Ruegeria* (*Rhodobacteraceae*) is a widely abundant bacterial symbiont of corals (Huggett and Apprill 2019) and was found enriched on the skin of the two fish species (*B. paula* and *P. labrovitattus*).

### Other bacterial classes enriched in fish skin microbiome

*Fusobacteria* were a third bacterial class enriched in the blennies’ skin microbiomes. In some cases, it constitutes a significant proportion of the microbiome (7–34%), showing a statistically higher relative abundance than in seawater (2–4%) or on substrates (0–6%). *Fusobacteria* were significantly more prevalent in *P. labrovittatus* (13%) than in *A. arnoldorum* (5%) and *B. paula* (2%). They were represented by a single order, *Fusobacteriales*, and four genera: *Cetobacterium* (0.74–11.59%), *Propionigenium* (1.23–1.59%), *Fusobacterium* (0–0.46%), and *Psychrilyobacter* (0.01–0.08%). *Cetobacterium* is one of the best-studied fish bacterial symbionts (Tsuchiya *et al*., 2008; Xie *et al*., 2022; Tsuchiya and Sugita, 2023; Zhao *et al*., 2023; M. Wang *et al*., 2025; S. Wang *et al*., 2025) commonly found in the gut and skin of many freshwater fish (Ramírez *et al*., 2018; Arias *et al*., 2019; Krotman *et al*., 2020; Coetzer *et al*., 2021) and occasionally on marine fish skin (Han *et al*., 2024). *Cetobacterium* isolates from the guts of aquatic animals can synthesize vitamin B12, helping their hosts meet vitamin requirements (Tsuchiya *et al*., 2008; Suzuki *et al*., 2022). Supplementing fish diets with *Cetobacterium* improves growth and overall health and enhances resistance to bacterial infections (Xie *et al*., 2022; Zhou *et al*., 2022; M. Wang *et al*., 2025; S. Wang *et al*., 2025). In the KEGG classification, vitamin B12 production was grouped with porphyrin metabolism, and *Cetobacterium* was predicted to be among the top-contributing genera to this functional category (Figure 5). *Cetobacterium* was also predicted to contribute the highest proportion (7.25%) to butanoate metabolism, whose primary product is butyric acid, a short-chain fatty acid found in the guts of omnivorous and herbivorous fish that has been shown to inhibit pathogens and provide health benefits to aquaculture fish (Yin et al. 2025). *Propionigenium* (*Fusobacteriales*), one of the most abundant genera in all three host species, is reported to be strictly anaerobic, succinate-fermenting, and to grow in marine sediments (von Hoyningen-Huene et al. 2022).

*Verrucomicrobiia* was another bacterial class with significantly higher relative abundance on fish skin (3–11%) than in seawater (0–1%) and substrate (0–4%). It was also significantly differentially abundant between the three fish hosts, comprising on average 1.4% in *B. paula,* 1.6% in *P. labrovittatus* and 6.4% in *A. arnoldorum. Verrucomicrobiia* has been only occasionally reported in fish skin, e.g. as a seasonal component in diadromous salmonids (Hamilton *et al*., 2023) and in lumpfish (Jacobsen *et al*., 2025). In our study, *Verrucomicrobiia* was represented by 62 ASVs, more than half (35) belonging to *Rubritalea*. Among the top 18 most abundant bacterial genera in the fish skin microbiome, *Rubritalea* accounted for an average of 2.8% total abundance. It is a key commensal on the skin of healthy European seabass and its abundance decreases following *Vibrio* infection (Cámara-Ruiz *et al*., 2021), which suggests *Rubritalea* may help maintain a healthy microbiome. Among the seven described *Rubritalea* species, five have been isolated from marine animals (Scheuermayer *et al*., 2006; Yoon *et al*., 2007, 2011) and in sponges it provides antioxidant protection by producing carotenoids and squalene (Shindo and Misawa, 2014), but its role in fish has not been studied yet. Our functional prediction suggested a role in the degradation of glycosaminoglycans (GAGs) and flavonoids. Since GAGs are major components of skin mucus (Tiralongo *et al*., 2020), and bacteria have GAG-degrading enzymes (Zhang *et al*., 2020), *Rubritalea* is most likely an important contributor to the maintenance and recycling of the fish mucosal surface. *Rubritalea* is significantly enriched in *A. arnoldorum* compared to the other two fish species, which may be related to the specific composition or thickness of its mucus coating. Two other genera within *Verrucomicrobiia* were enriched in fish skin: *Haloferula,* which is sensitive to salinity changes and reduces its abundance at lower salinities (Lai *et al*., 2022), and *Roseibacillus,* found on lumpfish skin (Jacobsen *et al*., 2025). Multiple *Roseibacillus* species have been isolated from algae, indicating that fish skin microbiomes can be temporarily colonized by epiphytic bacteria, particularly in species that spend time in algal-covered habitats.

The *Spirochaetia* class was also enriched in the fish skin microbiomes, being represented by 10 ASVs classified as *Treponema*. These are often parasites in a variety of animal hosts, including humans, but non-pathogenic species are also found in the oral cavities and intestinal tracts of humans and other animals (Miller et al., 1992; Edwards et al., 2003). *Treponema* has been detected in the gut of marine herbivorous fish living on rocky reefs in northern New Zealand and temperate Australia (Pisaniello et al., 2023). Interestingly, the presence of *Treponema* was restricted to mucosal surfaces in the gut (Pisaniello et al., 2023), suggesting that the mucus-covered skin of combtooth blennies may also provide a suitable habitat.

Within the class *Brevinematia, Brevinema* was among the most abundant genera in the skin microbiomes. It is more commonly found in animal guts, including Atlantic salmon (Li *et al*., 2021) and European seabass (Alfonso *et al*., 2023), but the thick mucus layer of combtooth blennies might provide a suitable habitat for what are normally intestinal microaerophilic bacteria, thanks to the presence of low-oxygen micro-niches.

Four additional bacterial classes were also enriched on fish skins: *Bacilli* (average contribution of 1.7%), *Balneolia* (0.6%), *Actinobacteria* (0.4%), and *Deferribacterota* (0.1%). These taxa make up part of the microbial “rare biosphere,” but they might have important roles in the microbiome, becoming beneficial (Lynch and Neufeld, 2015) or opportunistic pathogens (Doni *et al*., 2023) for the host under certain conditions. *Balneolia* and *Deferribacterota* are newly designated phyla (Oren and Garrity, 2021), which makes a comparison to older studies difficult. *Actinobacteria* and *Bacilli* (formerly *Firmicutes*) are consistently reported in fish skin microbiomes, but because of their low abundances, potential roles are rarely discussed (Larsen *et al*., 2013; Chiarello *et al*., 2018; Doane *et al*., 2020; Jacobsen *et al*., 2025). For example, (Tapia-Paniagua *et al*., 2018) found differences among low-abundance Bacilli between healthy fish skin, with higher prevalence of *Staphylococcus* and *Lactobacillus,* and skin lesions, enriched in *Streptococcus* and *Granulicatella*. In our data, five genera within *Bacilli* were enriched in fish skin: *Staphylococcus* (average relative abundance on fish skin 0.47%), *Turicibacter* (0.23%), *Streptococcus* (0.11%), *Mycoplasma* (0.5%), and *Lactobacillus* (0.003%). In addition, five *Actinobacteria* genera were enriched on the skin of blennies: *Cutibacterium* (0.11%), *Corynebacterium* (0.08%), *Mycobacterium* (0.06%), *Bifidobacterium* (0.02%), and *Actinomyces* (0.02%). Even though all the above genera contributed little to the total relative abundance, their functional profiles suggested they might play important roles in their host’s health through antimicrobial compound production and vitamin biosynthesis (Table S6). For example, *Corynebacterium* and *Mycobacterium* were predicted to contribute significant proportions to carbapenem biosynthesis, and *Staphylococcus* was one of the contributors to biotin (vitamin B7) synthesis.

### The skin of supratidal A. arnoldorum hosts more photosynthetic cyanobacteria

Differential abundance analysis of predicted functional categories identified several biochemical pathways that differed among the three fish host species. Specifically, photosynthesis was detected at higher levels in the skin microbiome of the supratidal *A. arnoldorum,* likely due to two cyanobacterial genera, *Phormidesmis* and *Schizothrix*, at higher relative abundances and contributing respectively 20% and 12% to photosynthesis. This could be explained by the greater light availability on supratidal rocks, which likely offers better conditions for oxygenic photosynthesis than the below-water environments. Similarly, the biosynthesis of carotenoids and other terpenoids (Figure 5) can explain the higher relative abundance of these cyanobacterial genera, and we speculate that these specialized metabolites could help protect the terrestrial *A. arnoldorum* from stress caused by prolonged light and UV exposure and oxidative stress (reactive oxygen species). Carbohydrate metabolism and specifically sulfoquinovose metabolism were also found at higher levels in the skin microbiome *of A. arnoldorum.* The utilization and degradation of sulfoquinovose, a sulfonated sugar derived from algal and cyanobacterial sulfolipids (Ma *et al*., 2025; Chen *et al*., 2026), was predicted to be mainly carried out by a variety of undescribed *Alphaproteobacteria* (18%), specifically a undescribed *Paracoccaceae* (12%), but also by *Vibrio* (10%) and *Granulosicoccus* (10%), which were more abundant in the skin microbiome of *A. arnoldorum*.

### The role of specialized metabolism and xenobiotic degradation

Many of the identified bacterial taxa enriched in fish skin microbiomes might be beneficial for the host thanks to their metabolic ability to produce specialized metabolites and vitamins such as carotenoids and/or terpenoids (*Paracoccacea UG, Schizothrix, Phormidesmis, Rubritalea*), vitamin B12 (*Paracoccacea UG, Cetobacterium*), vitamin B6, folate, and lipoid acid (*Alteromonas*), biotin (*Cetobacterium, Schizothrix*), and panthotenate and nicotinamide (*Granulosicoccus*). Furthermore, various genera (*Acinetobacter*, *Pacoccacea UG*, *Granulosicoccus*, *Alteromonas*, *Kordiimonas*, *Cetobacterium*, *Aranicella*, *Burkholderiales* UG) might have the metabolic potential to synthesize antimicrobial metabolites (Figure 5) that may help maintain healthy skin microbiomes.

Several other bacterial taxa enriched in the fish skin microbiomes seem to possess the ability to degrade various xenobiotics were enriched in the subtidal *B. paula*. Pathways for the degradation of toluene, nitrotuene, and xylene were higher in *B. paula* and *P. labrovitattus* in comparison to *A. arnoldorum.* The most abundant genera predicted to contribute the highest proportions to xenobiotics degradation were *Acinetobacter*, *Paracoccaceae* UG, and *Granulosicoccus, Alphaproteobacteria UG, Cetobacterium* and *Parendozoicomonas*. Bisphenol and PAHs are persistent pollutants that accumulate in marine sediments and are thus particularly toxic to bottom-dwelling organisms. In marine environments, abiotic degradation of these xenobiotics is limited and microbial metabolism is the primary removal pathway. The higher importance of these pathways in submerged (subtidal and intertidal) fish may reflect their continuous exposure to these pollutants compared to the supratidal *A. arnoldorum.* In addition, some of these chemicals, including xylene, can be degraded by sunlight, which is more intense out of water, and thus might explain the absence of xylene-degrading bacteria on *A. arnoldorum* skin.

### Bacterial motility in the mucus layer of combtooth blenny skins

Several bacterial taxa enriched in fish skins appear to be adapted to live in the mucus layer of combtooth blennies. This layer is secreted by superficial epithelial and goblet cells and consists of sulfated acid glycoproteins and mucus (Whitear and Mittal 1984). This mucus not only provides a viscous environment but also probably low-oxygen micro-niches for microaerophilic bacteria typical of the gut. Some bacteria enriched in fish skin (*Rubritalea* and *Cetobacterium*) have the potential to degrade glycosaminoglycans, which are constituent components of mucus and the extracellular matrix, as well as other glycans (*Photobacterium* and *Rubritalea*). The significant contribution to amino sugar and galactose metabolism observed in *Vibrio* and *Photobacterium* and the enrichment in amino acid degradation pathways in *Cetobacterium* (among which lysine, 10.5%) could also correlate with their capacity to degrade and utilize mucus and extracellular matrix components. Notably, 15 of the 22 genera shown in Figure 5 (i.e., all except *Dokdonia, Phormidesmis, Cetobacterium, Arnicella, Rubritalea, Propionigenium, Lentisphaera*, and undescribed genera—UG) contain motile species, an important ability to navigate a viscous environment such as the blenny skin mucus layer. Moreover, some bacteria present cell morphologies that are vibrioid (in *Vibrio* and *Marivibrio*; members of *Gammaproteobacteria, Enterobacterales*) and/or spiral forms (*Marivibrio* and *Brevinema*), which are well-known adaptations for motility in viscous environments and might be important for virulence in pathogenic bacteria (Wadhwa and Berg 2022; Bartlett et al. 2017; Xu, Koizumi, and Nakamura 2020).

## Conclusions

The skin microbiomes of three combtooth blenny species were studied in the context of adaptation to increasingly amphibious lifestyles. Skin-associated communities were distinct from those on surrounding seawater and rock substrates, indicating that fish skin is an important ecological filter. A second ecological filter is imposed by the intertidal gradient, as evidenced by fish skin microbiomes that diverge progressively with the increasing terrestriality of host fish, the supratidal amphibious *Alticus arnoldorum* harboring the most distinct skin microbiome. *Pseudomonadota* (*Alpha and Gammaproteobacteria*) dominated the skin microbiome and the balance between *Alpha*- and *Gammaproteobacteria* shifted with habitat: *Alphaproteobacteria* were more abundant in the subtidal *B. paula* and intertidal *P. labrovitattus*, whereas *Gammaproteobacteria* were enriched in the supratidal *A. arnoldorum.* Skin communities also showed enrichment of epiphytic-associated taxa, consistent with benthic lifestyles and frequent contact with algae-covered substrates. Functional predictions suggest that the skin microbiomes of these fish differ not only in taxonomic composition but also in metabolic potential. Pathways related to algal-derived compound metabolism (e.g., photosynthesis and sulfoquinovose-associated carbohydrate metabolism) were comparatively enriched in *A. arnoldorum*, consistent with greater exposure to light in supratidal rocks. In contrast, subtidal and intertidal species showed a higher predicted representation of xenobiotic degradation pathways, which may be explained by their higher exposure to dissolved and sediment-associated contaminants in underwater environments. While these functional inferences require validation using metagenomics, metatranscriptomics, and culture-based experiments, they provide testable hypotheses that link habitat, microbial community structure, and functional potential across the intertidal ecological gradient. Our study also emphasized the unique nature of the scaleless mucus-covered skin of combtooth blennies. The presence of several anaerobic or microaerobic taxa, typically associated with the gut (e.g., *Cetobacterium*), suggests that the thick mucus layer likely provides low-oxygen micro-habitats for these species. Moreover, many of the detected bacterial taxa are motile or have vibrioid or spiral cell morphologies as adaptations to viscous environments. This is the first study to assess how fish skin microbiomes vary across species with increasingly terrestrial lifestyles, positioning the intertidal as a natural experiment in microbiome assembly under variable exposures to water, oxygen, and environmental chemicals. Future work should resolve taxa at strain level and test functional mechanisms—such as vitamin and micronutrient synthesis, antioxidant production, mucin/glycan utilization, and xenobiotic transformation—to clarify when dominant skin microbes act as commensals, mutualists, or opportunists under changing environmental conditions.

## Data Availability

Raw sequencing data have been deposited in the NCBI Sequence Read Archive (SRA) under BioProject accession PRJNA1459627. Associated metadata and processed data are available at https://github.com/ewrubin/combtooth-blenny-microbiome

## Funding

This project was funded by the Spanish State Research Agency (AEI) MICIU/AEI/10.13039/501100011033, ESF+ and ERDF/EU (Grants RyC2022-038245-I,

PID2023-152168NB-I00, and CNS2025-165952) and the Leibniz Institute for the Analysis of Biodiversity Change (LIB), Germany.

## Acknowledgements

We are grateful to Elizabeth A. Surovic, Héctor Torrado, and David Combosch for their support during fieldwork, and to Sarah Lemer and Héloïse L. Rouzé for facilitating laboratory work. We thank Jeffrey Quitugua, Brent Tibbatts, William Paulino II, and the remaining staff from Government of Guam Department of Agriculture, Division of Aquatic & Wildlife Resources (DAWR) for collection and export permits (SCR-23-006) and for their continuous support during fieldwork in Guam. We also thank the University of Guam UOG-IACUC committee and the University of New South Wales Animal Care and Ethics Committee for ethical permits. We are grateful to Laurie Raymundo for support in obtaining sampling permits and endorsing our stay at the University of Guam Marine Laboratory. We thank Angela Duenas, the Biorepository team, and the rest of the University of Guam Marine Laboratory Staff for supporting our visit.

