## Supplementary figures and images for "Skin microbiome mirrors habitat divergence in amphibious combtooth blenny fish (Teleostei, Blenniidae)"

### Figure S1

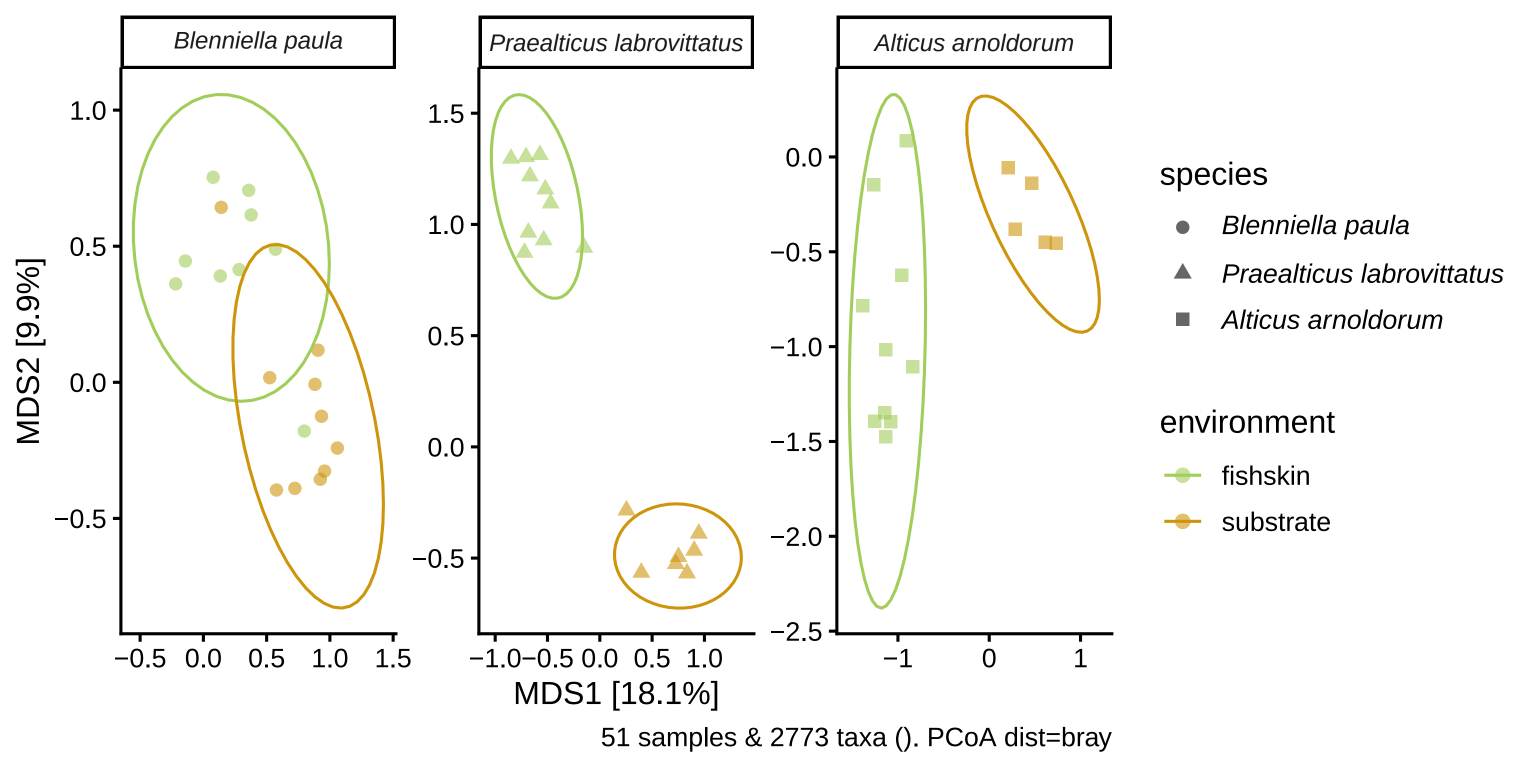

### Figure S2

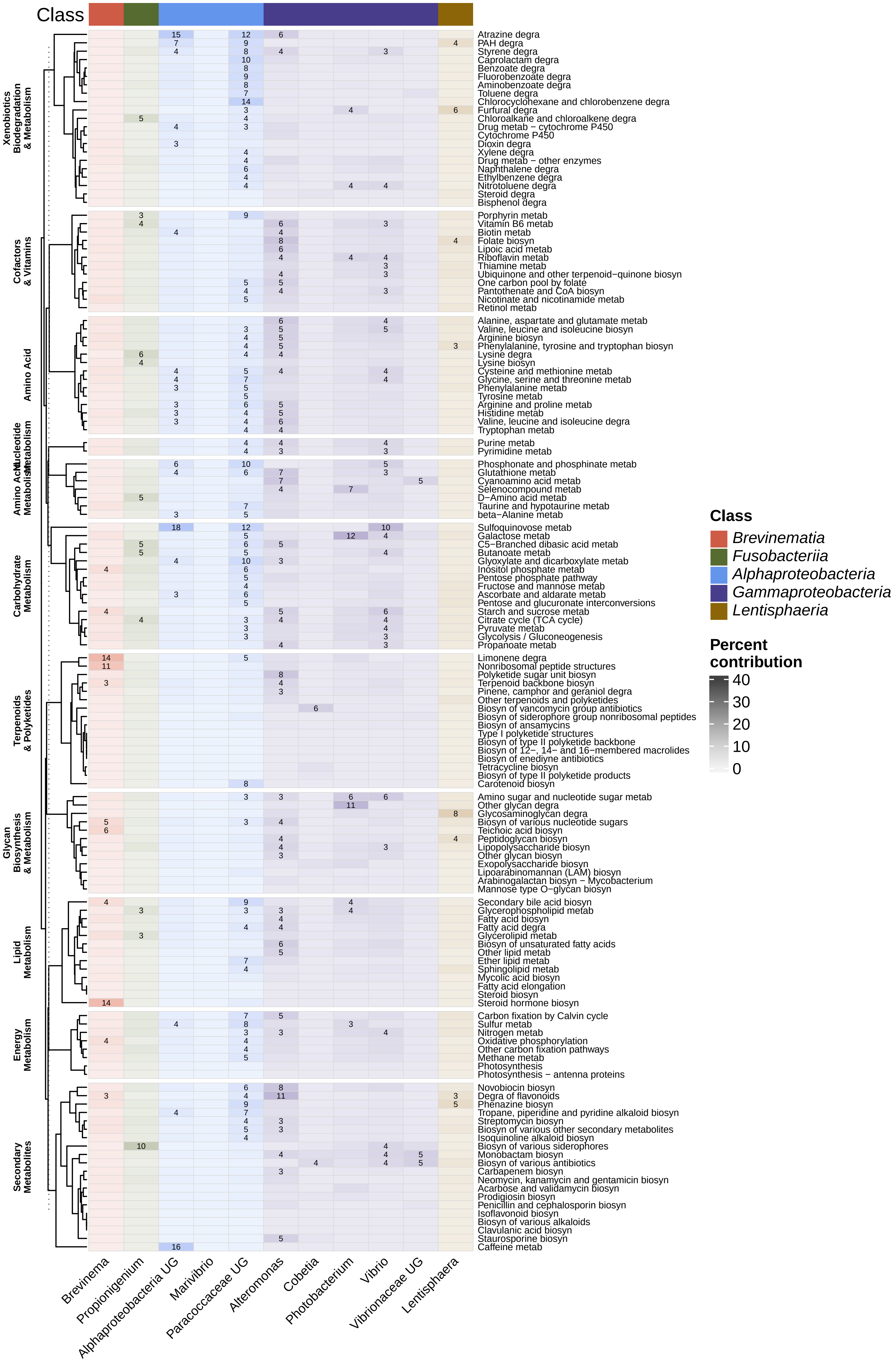
